# Rationale design of peptide inhibitors for *α*-Synuclein liquid condensates and fibrillar aggregates using multiscale modelling approach

**DOI:** 10.1101/2025.01.12.632580

**Authors:** Srinivasan Ekambaram, Santosh Prajapati, Anand Srivastava

**Affiliations:** Molecular Biophysics Unit, Indian Institute of Science, C. V. Raman Road, Bangalore, Karnataka, India 560012

## Abstract

*α*-Synuclein is an intrinsically disordered protein (IDP) whose aggregation is implicated in Parkinson’s disorder. Herein, we computationally design a *α*-Synuclein derived potential peptide inhibitor against the protein’s monomeric, fibrillar and liquid condensate forms using multi-scale molecular modelling approaches. Since the conventional structure-based design paradigm often is not applicable to these highly labile IDPs, we first develop a pipeline to generate an exhaustive library of small candidate peptides from an available repository of *α*-Synuclein 70 *µ*s all-atom molecular dynamics (AAMD) trajectory data. We then use high throughput screening techniques such as PATCHDOCK and HPEPDOCK as well as AAMD simulations to arrive at a single candidate peptide. AAMD simulations data show that *α*-Synuclein bound peptide chain leads to an expanded conformational ensemble for the chain and also reduces the *β*-sheet propensity of the protein’s fibrillar amylogenic aggregates. Coarse-grained simulations using HPS-Cation forcefield with 100 chains of *α*-Synuclein and varying levels of candidate peptides shows decreased density and increased apparent critical temperature of the condensate system. Our detailed molecular interactions analyses show that peptides bind to the *α*-Synucleins through the “dynamic shuttling mechanism” where interaction are frequently made and broken around a given set of structurally proximal residues, which likely softens the dynamic interaction network in the condensates. Together, we could illustrate the inhibitory effect of the final designed peptide against distinct forms of *α*-Synuclein monomer and aggregates. Our work provides a multiscale simulation based prescription towards the futuristic development of therapeutic strategies against the disordered proteins.

## 1 Introduction

*α*-Synuclein is an aggregation-prone intrinsically disordered protein (IDP) that plays a prominent role in the pathogenesis of Parkinson Disease (PD) and other synucleinopathies. ^1–6^

Structurally, *α*-Synuclein is a 140 amino acids (AA) long IDP with a N-terminal amphipathic region (residues 1-60) and a hydrophobic rich non-amyloid *β* component (NAC) region (residues 61-95), which can drive aggregation and formation of cross-*β*-structures.^7–9^ The C-terminal disordered region (residues 96-140) is rich in Aspartic and Glutamic acids and largely exist as a random coil structure, which is implicated in nuclear localization and interactions with small molecules.^10,11^ Reports from various studies have confirmed that *α*-Synuclein exist as an ensemble of conformations in solution. ^7,12^ In contrast, the recently resolved cyro-EM model has revealed that in the fibril form, the *α*-Synuclein takes up the *β*-sheet structure that is considered toxic to neuronal cells.^13^ Though the exact molecular mechanisms leading to formation of the pathological fibrillar structure in IDPs is still elusive, several studies have shown the ability of many IDPs to experience liquid-liquid phase separation (LLPS) that can eventually solidify into gel-like structures or amyloid-like aggregates.^14–18^

IDPs are found to be common pathological factors in various human diseases such as cardiovascular, diabetes and neurological disorders such as such as Alzheimer’s, Parkinson’s (PD) and amyotrophic lateral sclerosis(ALS). ^1–3^ Among the several line of therapeutic interventions followed in IDP-related cure, small drug-like molecules inhibitors are attractive candidates. However, conventional drug design strategies that are applied for folded proteins with well-defined binding sites and pockets can not be used for IDPs. This is because IDPs such as *α*-Synuclein lack a reference structure and exist as an ensemble of conformations with frequent sub-microsecond transition between the various states. As such IDPs are often considered undruggable. ^19–21^ Targeting the aggregation process of the IDPs associated with disease has emerged a primary route for drug discovery in the area, and a range of inhibitors for A*β*, Tau, and *α*-synuclein aggregation have been tried in the literature.^22–32^ Numerous studies, both computational and experimental, have stipulated that small molecules could act as inhibitors against the *α*-Synuclein.^11,33–39^ However, in the absence of successful compounds at the clinical stage, there is a pressing need to develop new strategies for the identification of more effective inhibitors of the aggregation process of these proteins.

Peptide-based inhibitors are an attractive alternative for IDPs as effective pharmacological agents in the development of modern medicine over the past decade. ^40,41^ Various peptide-based inhibitors have been studied against *α* Synuclein.^42–45^ Here, we focus on the design of a new therapeutic peptide derived from the *α*-Synuclein sequence that acts as an inhibitor against it. *α*-Synuclein-derived peptides offer several promising attributes as a therapeutic strategy. These peptides can directly engage with aggregation-prone regions of *α*-Synuclein and potentially disrupt its propensity to misfold and form aggregates. Owing to their intrinsic origin, such peptides exhibit high specificity and affinity for *α*-Synuclein and minimize off-target effects. Moreover, the relatively small size of peptides facilitates cellular penetration and enhances their accessibility to intracellular *α*-Synuclein. By competing with full-length *α*-Synuclein for binding partners or aggregation nucleation sites, these peptides may effectively inhibit the formation of toxic oligomers and fibrils.

Multiscale computational approaches combining molecular docking and moleculr dynamics (MD) simulations has the potential to emerged as a powerful strategy for studying protein aggregation inhibition using peptides. In this pipeline, initial high-throughput screening of potential peptide inhibitors is performed using molecular docking, which rapidly evaluates their binding poses and affinities. Promising candidates identified through docking are refined through MD simulations allowing for the assessment of peptide-protein interactions in a dynamic, fully-solvated environment to assess electrostatic and hydrophobic contributions to binding energy. This method allows for a more realistic evaluation of protein-ligand interactions in a dynamic environment. Several studies have employed a multiscale computational approach combining molecular docking and MD simulations to optimize drug design and selection. ^46–48^ Such methods provide insights into the stability of peptide-protein complexes, conformational changes, and potential mechanisms of aggregation inhibition. By bridging the gap between static docking poses and dynamic protein-peptide interactions, this multiscale approach offers a more comprehensive understanding of aggregation inhibition processes, potentially accelerating the discovery of novel therapeutic peptides. However, this framework is challenging to implement for IDPs due to their conformational flexibility. In this work, we have enhanced this framework such that it could be used to design effective peptide inhibitors for labile IDPs.

The rest of the paper is organized as follows. This introduction section is followed by the Results and Discussion section where we report the pipeline we have created to generate the peptide library, screen for candidate peptides and design the final effective peptide for simulation trials on monomeric, fibrillar and condensate form of *α* Synuclein. We analyse these systems and report our findings related to effect of peptide on protein behavior. We summarize our findings in the Conclusion section and also highlight the scope and limitations of the prescription. We provide details of the systems under considerations and simulations methods in the Material and Methods section. All input files needed to initiate molecular simulations and full trajectory data of all simulations for all systems considered in this work are publicly available on our laboratory server for download. The server data can be accessed via our laboratory GitHub link: codesrivastavalab/synuclein-KFQVT.

## 2 Results and Discussion

The first step in our workflow is to generate an exhaustive library of candidate peptides from the conformation ensemble of *α*-synuclein. In the following sections, we discuss how the peptide library is screened for the final candidate peptide. Once we arrive at the most suitable peptide candidate, we carry out large scale all-atom molecular dynamics (AAMD) simulations to explore the changes in conformational landscape of *α*-synuclein due to the peptide. We also carry out AAMD on effect of peptide on the fibril structure of *α*-synuclein. Lastly, using coarse-grained molecular dynamics (CGMD) simulations, we look at effect of the candidate peptide on the liquid condensate properties of *α*-synuclein. Broadly, our results show that the candidate peptide has a high residence time with *α*-synuclein molecule and it changes the conformations landscape of the protein – both at the single molecule level and the fibrillar structure. The peptide also decreases the density and the “apparent” critical temperature of the liquid condensate. Below, we discuss these aspects and our findings in greater details.

### 2.1 Generating the candidate peptide library

We initially obtained the 73-microsecond AAMD simulation data of *α*-Synuclein from Paul Robustelli and co-workers^49^ to design a peptide inhibitor. To identify representative conformations of *α*-synuclein, t-distributed stochastic neighbor embedding (t-SNE) based IDP-clustering algorithm, recently developed in our group,^50^ was employed to analyze the conformational ensemble generated through molecular dynamics simulations. The clustering analysis yielded 50 distinct conformational clusters (represented as 50 different groups in the t-SNE projection shown in Fig. 1A). We report the respective probabilities of each conformer in Fig. 1B where the population of each conformation is between ∼ 1 % to ∼ 3.5 %. From this analysis, we selected the top conformations based on two criteria: high probability and location within dense clusters in the t-SNE projection. This approach led to the identification of 14 representative conformations of *α*-synuclein, designated as AS-1, 6, 8, 9, 10, 12, 13, 19, 21, 24, 28, 38, 41, and 48 (represented in Fig. 1A with arrows showing the central conformation of these clusters). These conformations were selected based on their high probability and location within dense clusters in the t-SNE projection, ensuring a comprehensive representation of *α*-synuclein’s structural diversity. Notably, these selected conformations accounted for 36% of the entire *α*-synuclein conformational population, providing a robust foundation for further analysis and peptide library development. The 14 representative conformations served as initial templates for the development of a peptide library, with the strategy for library generation illustrated in Figure 1C. This approach involved systematically deriving peptide sequences from different regions of each *α*-synuclein conformation, taking into account various structural features and potential interaction sites. By employing this method, we generated an average of 1,900 peptides per conformational library, resulting in a total library size of approximately 13,230 peptides. This extensive library encompasses a wide range of peptide lengths, sequences, and structural propensities, reflecting the conformational plasticity of *α*-synuclein. A key feature of this study is the derivation of peptides from these diverse conformations, which provides an extensive array of peptide structures with varying secondary structural elements. This diversity is particularly significant given *α*-synuclein’s nature as an intrinsically disordered protein (IDP). The peptide library captures the full spectrum of structural possibilities of a given sequence stretch of *α*-synuclein, including *α*-helical, *β*-sheet, and random coil conformations, as well as various intermediate states. This structural heterogeneity is crucial for exploring potential inhibitory mechanisms against *α*-synuclein aggregation, as it allows for the identification of peptides that may interact with different conformational states of the protein throughout its aggregation pathway. Furthermore, the large size and diversity of the peptide library increase the likelihood of identifying highly specific and potent inhibitors of *α*-synuclein aggregation. By sampling a broad conformational space, we enhance our chances of discovering peptides that can effectively target key regions involved in *α*-synuclein oligomerization and fibril formation. This approach also allows for the exploration of structure-activity relationships, potentially leading to the identification of critical structural features necessary for inhibitory activity. To illustrate this structural diversity, we examined the peptide GVAEAAGKT, which exhibited multiple secondary structure features, including *α*-helix, coil, and turn conformations, as depicted in Fig. 1D. This observation supports our hypothesis that the developed peptide library encompasses a wide range of conformations with distinct secondary structure features, a crucial factor in the design of effective peptide-based inhibitors. In our library design, we deliberately constrained peptide lengths to a range from 3 to 9 amino acids. This decision was informed by previous studies indicating that shorter peptides often demonstrate superior potency as inhibitors, exhibiting improved bio-availability and enhanced resistance to proteasomal degradation. Furthermore, the extensive conformational sampling is crucial for accurate docking studies, enabling a thorough investigation of potential binding modes and interactions with *α*-synuclein. By considering the dynamic nature of *α*-synuclein aggregation, our approach increases the likelihood of identifying effective inhibitors capable of adapting to various protein conformations. By adopting this strategy, we aimed to identify peptide inhibitors capable of effectively modulating the aggregation propensity of *α*-synuclein through disruption of key intermolecular interactions. Thereby, employing t-SNE clustering to capture the conformational heterogeneity of *α*-synuclein, we have developed a robust approach for generating a diverse peptide library. This methodology provides a solid foundation for the rational design of novel peptide-based inhibitors targeting this intrinsically disordered protein. Our strategy prioritizes the targeting of prevalent *α*-synuclein conformers, which we hypothesize will enhance the likelihood of developing effective inhibitors.

**Figure 1:**
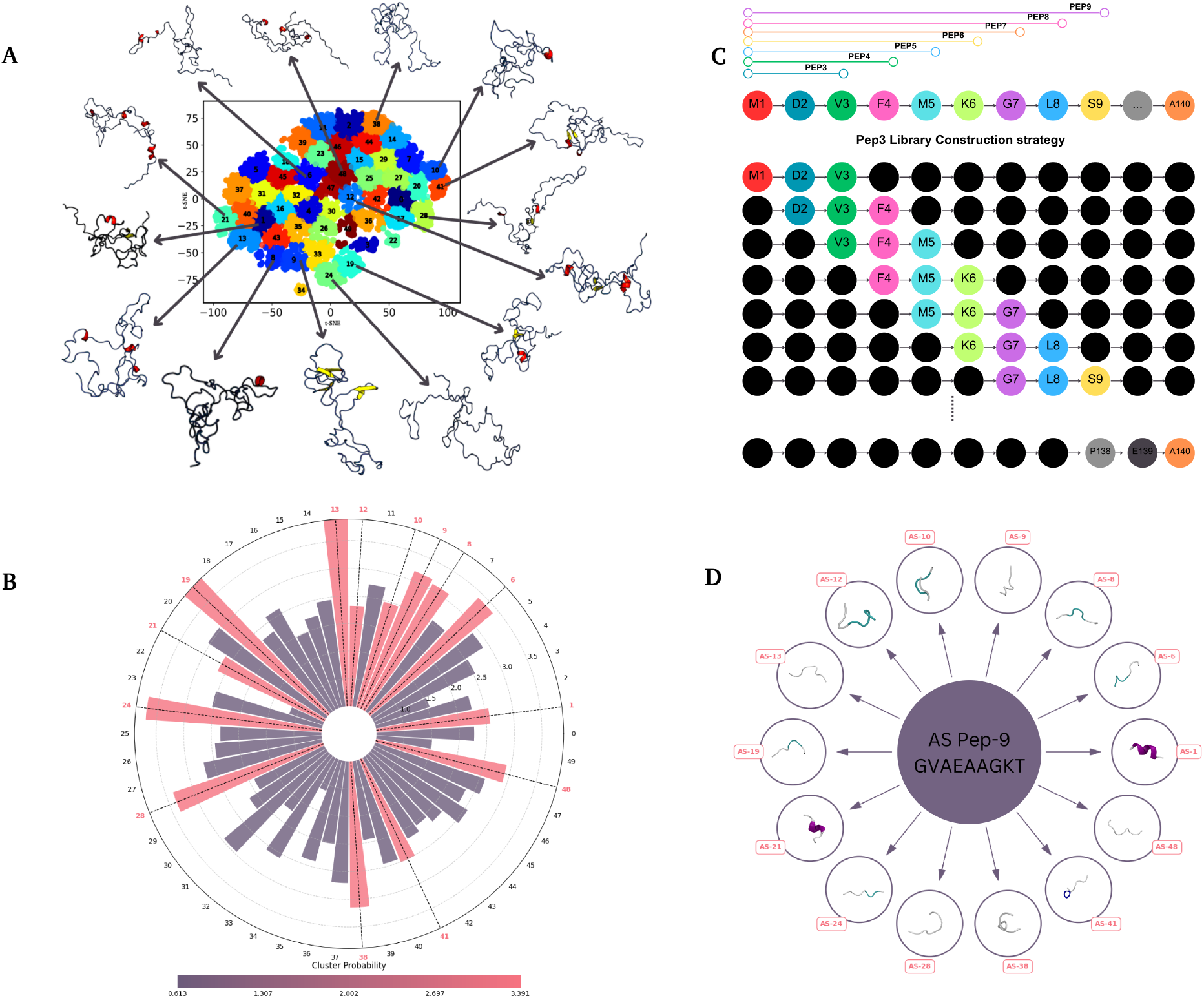
(A) t-SNE clustering of 73-microsecond simulation of *α*-Synuclein conformations. We have selected the 14 different conformations from the densely populated cluster with distinct conformations for developing the peptide library in this study. (B) The cluster probability of the A-Synuclein conformations from t-SNE clustering with the most probable cluster used in the study is highlighted in pink. (C) The peptide library building strategy used in this study to construct peptide libraries from 3mer to 9mers. (D) The secondary structure comparison of the same sequence attains multiple secondary structure stores, which shows the intricate properties of the IDP peptides.

### 2.2 High throughput screening pipeline suggests 7-mer sequence of TQKEQVT from ***α***-Synuclein as best peptide candidate

The peptide library builder algorithm provided us with 13,230 different peptides structures with varying peptide lengths from 3-mer to 9-mers (Fig. 2). For docking studies, we treat the *α*-Synuclein conformations as the substrate (Fig. 2(a)) and peptides as ligands (Fig. 2(b)). Fig. 2(c) is the schematics of our pipeline. We used PATCHDOCK^51^ for the initial screening of peptides since its algorithm works by molecular shape complementarity. The top binding peptide from each subset of the peptide library was chosen for further refinement in the docking with HPEPDOCK.^52^ We initially employed PatchDock to generate complementary binding poses for the protein-peptide complexes. These poses were subsequently evaluated and refined using PRODIGY.^53^ In the Fig. 2(d), we report the energy scores calculated by Prodigy for PatchDock conformations and also the scores from HPEPDOCK for top candidate peptides selected from PatchDock conformations. In the figure, top candidate peptides are shown in Orange cartoon with sticks. The highest-scoring peptides from each library were further refined through re-docking with HPEPDOCK, which offers superior accuracy in protein-peptide binding predictions compared to other available methods.^54^ The seven top peptides in each library have better affinity for the *α*-synuclein conformations computed via PRODIGY. The seven peptides are: QKT, QVTN, EQVTN, KEQVTN, TKEQVTN, KKDQLGKGN, and KTKEQVTNV. The top peptides were further subjected to HPEPDOCK for accurate prediction of the protein peptide pose with affinity score. The results from the rigid docking study was followed by AAMD simulations.

**Figure 2:**
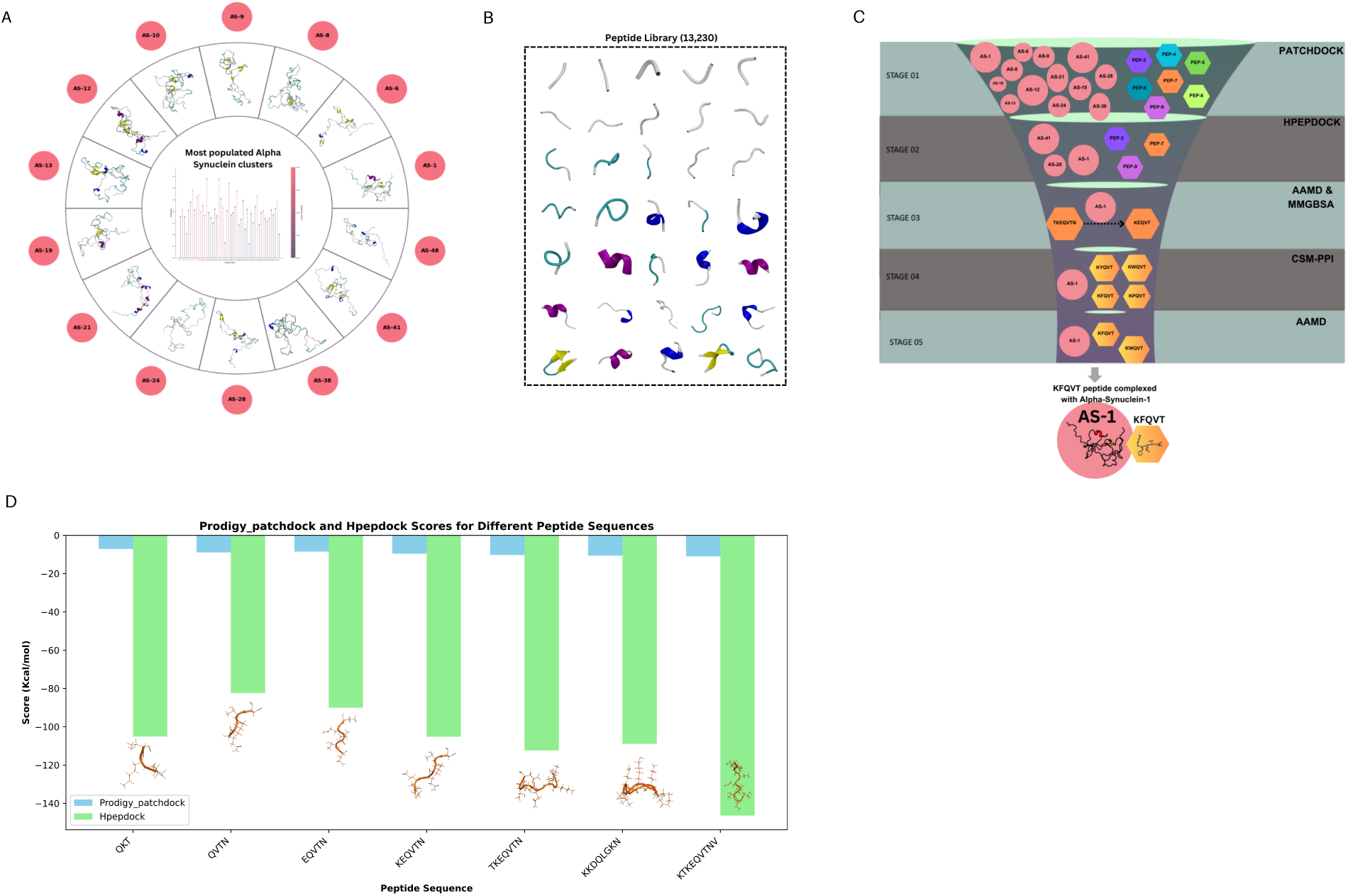
(a)The most populated *α*-synuclein conformers clusters selected from this study. (b) Image demonstrating the distinct variations in the secondary structural features of *α*-synuclein. (c) Our pipeline for screening of inhibitor peptide from the created repository (d) Energies calculated by Prodigy for PatchDock conformations and scores from HPEPDOCK for top candidate peptides selected from PatchDock conformations. Top candidate peptides are shown in Orange cartoon with sticks.

For the AAMD simulations, we monitored each complex for 100 ns and screened out the peptides that detached from the *α*-Synuclein within that timescale. For peptides that stayed bound for 100 ns, we extended the runs for 400 ns. Finally, a 7-mer peptide TKEQVTN showed consistent binding with one of the conformations of *α*-Synuclein chain for the duration of the run. We chose TKEQVTN as a base model peptide for the peptide design study with the objective to modify the sequence and length around the TKEQVTN sequence to arrive at the final candidate peptide.

### 2.3 Simulations predicts KFQVT as an effective ligand for ***α***-Synuclein

During the peptide design, we carefully evaluated the amino acids in the peptide that contributed to the binding with *α*-Synuclein chain using gmxMMPBSA calculations. ^55^ From the gmxMMPBSA results, we could decipher that the KEQVT amino acids in the TKEQVTN peptide showed higher energy contribution and hence we chose KEVQT for the next design phase (Fig. 3(a,b)). Further, we used the CSM-PPI web server^56^ to explore if we could design a modified peptide with higher binding affinity than KEVQT. The algorithm takes the peptide-substrate complex and carries out mutations on the initial screened peptide residues (KEQVT in our case) to arrive at an alternate peptide with higher binding with the provided substrate. For the KEQVT-*α*S complex, our results showed four different mutant peptides with higher affinity: KWQVT, KFQVT, KPQVT and KYQVT. For these four alternative peptides, we performed docking using HPEPDOCK and computed the binding energy values using HWAKDOCK.^57^ Docking results showed that the peptides KFQVT and KWQVT showed better binding affinity than others. HPEPDOCK and MMGBSA scores for KEQVT was found to be −94.43 and −32.97, respectively. We denoted that score as KEQVT(−94.43, −32.97) here. The other variants were found to have the following scores: KWQVT(−129.64, −42.03), KPQVT(−116.81, −31.96), KFQVT (−132.63, −43.39) and KYQVT(−123.35, −33.54). KWQVT and KFQVT are two most promising candidates after this screening. We followed this up with AAMD simulation protocol as discussed in the previous paragraph and found that the KFQVT peptide had better binding than KWQVT with *α*-synuclein chain. Further, we carried out a microsecond long AAMD simulation with KFQVT-*α*S and the complex was stable for 95% of the simulation time. Please see the corresponding movie files for KFQVT-*α*Synuclein complex simulation (movieS1a-movieS1d.mpeg) in our supplenementary information (SI). The description for each of the movie file is available in the SI pdf file. The link to the trajectory location can be found in the Github repository.

**Figure 3:**
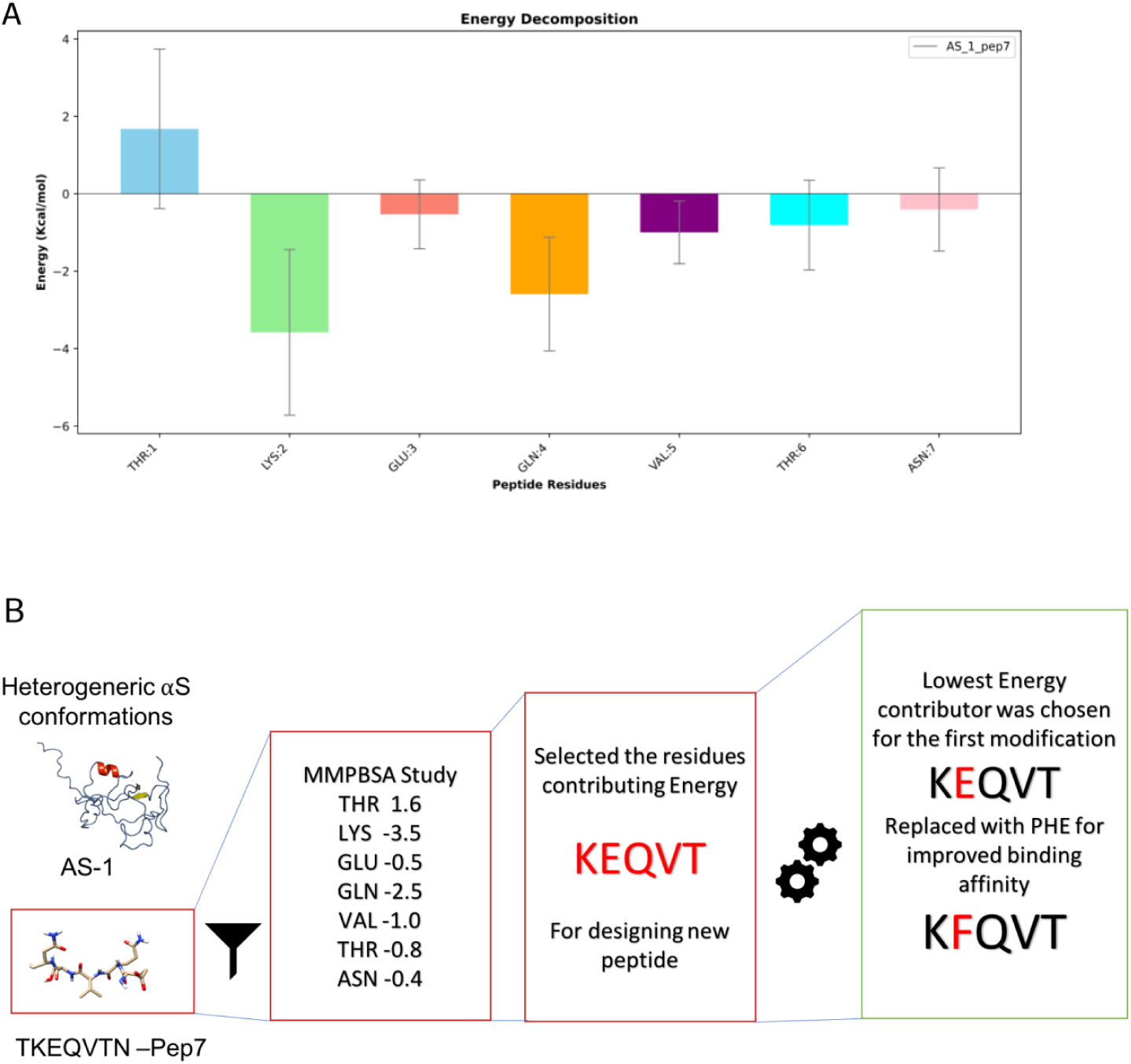
(a) The streamlined methodology for choosing the KFQVT peptide. The initial screening of the peptides through computational docking suggested towards the pep7 (TKEQVTN) as the top candidate for the peptide inhibitor having better residence time with the *α*-Synuclein over the MD simulation. The gmxMMPBSA analysis aided in elucidating the per residue energy contribution of the peptide towards the binding interaction with *α*-Synuclein protein. Thus, we truncated the pep7 by selecting KEQVT for further peptide design study. We utilised the CSM-PPI webserver to identify the mutations that could aid the augmenting the binding affinity of the peptide with the protein. (b) Per-residue energy decomposition analysis of Pep7 bound to AS1, calculated using gmxMMPBSA averaged over a 300 ns MD simulation. The peptide remained fully bound to the protein throughout this simulations.

To confirm that the KFQVT binds stably with *α*-Synuclein for extended time, we ran multiple replicates with different starting conformation of *α*-Synuclein. From the overall 5 *µ*second AAMD runs, the peptide was found to have a stable interaction with *α*-Synuclein for greater than 4 *µ*seconds. This exhibits reasonable consistent binding of the peptide against the *α*-Synuclein. Next, we compared the effect of peptide on the conformational landscape of *α*-Synuclein.

### 2.4 KFQVT disrupts the conformational landscape of ***α***-Synuclein

We computed the compactness and residual flexibility of the peptide-*α*S complex (Fig. 4A) and compared the results against the APO monomeric state of the protein using AAMD simulations. The peptide bound *α*-Synuclein exhibited an enhanced average solvent accessible surface area (SASA) (see Fig. 4B) as compared to the APO state. Also, the end to end distance (R*_ee_*) and radius of gyration (R*_g_*(Fig. 4C and Fig. 4D, respectively) showed an extended state of the protein when bound to the peptide. In general, binding to another molecules results in geometric compactions due to the enhanced enthalpic interactions between the partner molecules. However, our data suggests that *α*-Synuclein experiences entropic expansion upon binding. Interestingly, at the interactions level, we do see a loss in the number of hydrogen bonds within the *α*-Synuclein monomer (see Fig. 4E). Our observation follows the trend from previous studies from Michele Vendruscolo’s group that reported the effect of small molecule inhibitors on the disordered proteins leading towards the entropic expansion resulting from an effective increase in the conformational entropy of the proteins.^58,59^ Similar observations were also made in two other recent AAMD simulation work on binding of small molecule Fasudil with *α*-Synuclein resulting in an expanded conformation of the monomeric complex^11^ and inhibition of stable dimeric contacts formation due emergence of aggregation-resistant long-range interactions. ^60^

**Figure 4:**
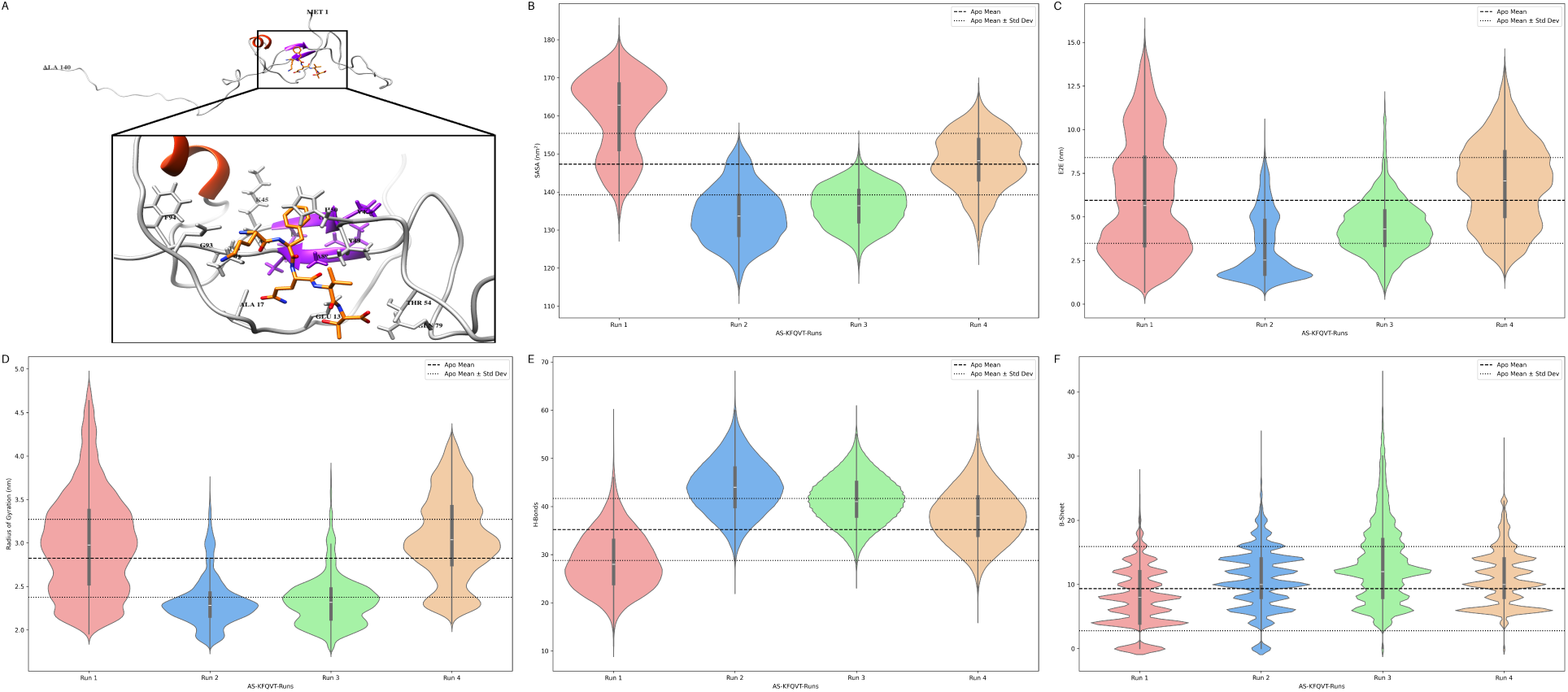
Geometrical assessment of the trajectories in apo and bound forms plotted across multiple runs. (a) *α*-Synuclein residues interacting with the derived peptide KFQVT (b) Solvent accessible surface area (SASA) showed that the system is more exposed to solvent in the complex state. (c) The end-to-end distance of the *α*-Synuclein was significantly increased upon the binding of the peptide. (d) The compactness of the protein adopted extended conformation in the bound state relative to the apo state. (e) Intramolecular hydrogen bonds were noticeably reduced when the peptide interacted with the *α*-Synuclein. (f) The propensity of the *β*-Sheets in *α*-Synuclein computed from the trajectory using DSSP in both apo and bound forms.

### 2.5 KFQVT disrupts residue-wise contacts in the NAC region

We computed the per residue secondary structural tendencies using the DSSP algorithm^61^ for both the APO and KFQVT-bound state of *α*-Synuclein to substantiate the conformational changes further. From the outcomes, we could infer that the beta-sheet propensity in the APO state was noticeably reduced upon binding with the peptide (Fig. 4F). Concomitantly, the propensity of the coil in the bound form was augmented compared to the apo state (54%). We also analysed the secondary structural changes over the residues in both the APO and bound form. As shown in Fig. 5A, probability of beta-sheets was reduced upon binding with pentapeptide – specifically at residual positions of NAC domain 70-75 and 85-90. The pentapeptide binding seems to lower the beta-sheet formation mostly in aggregation-prone regions reported from the experimental studies. Also, the most abundant formations of coils with a probability greater than 0.75 occur in the bound state at positions 60-70, 85-90 and 95-100. The bound complex shows an increased likelihood of the coil content with noticeable loss in beta-sheet probability, which could explain the augmented flexibility and shifted conformational propensities described above.

**Figure 5:**
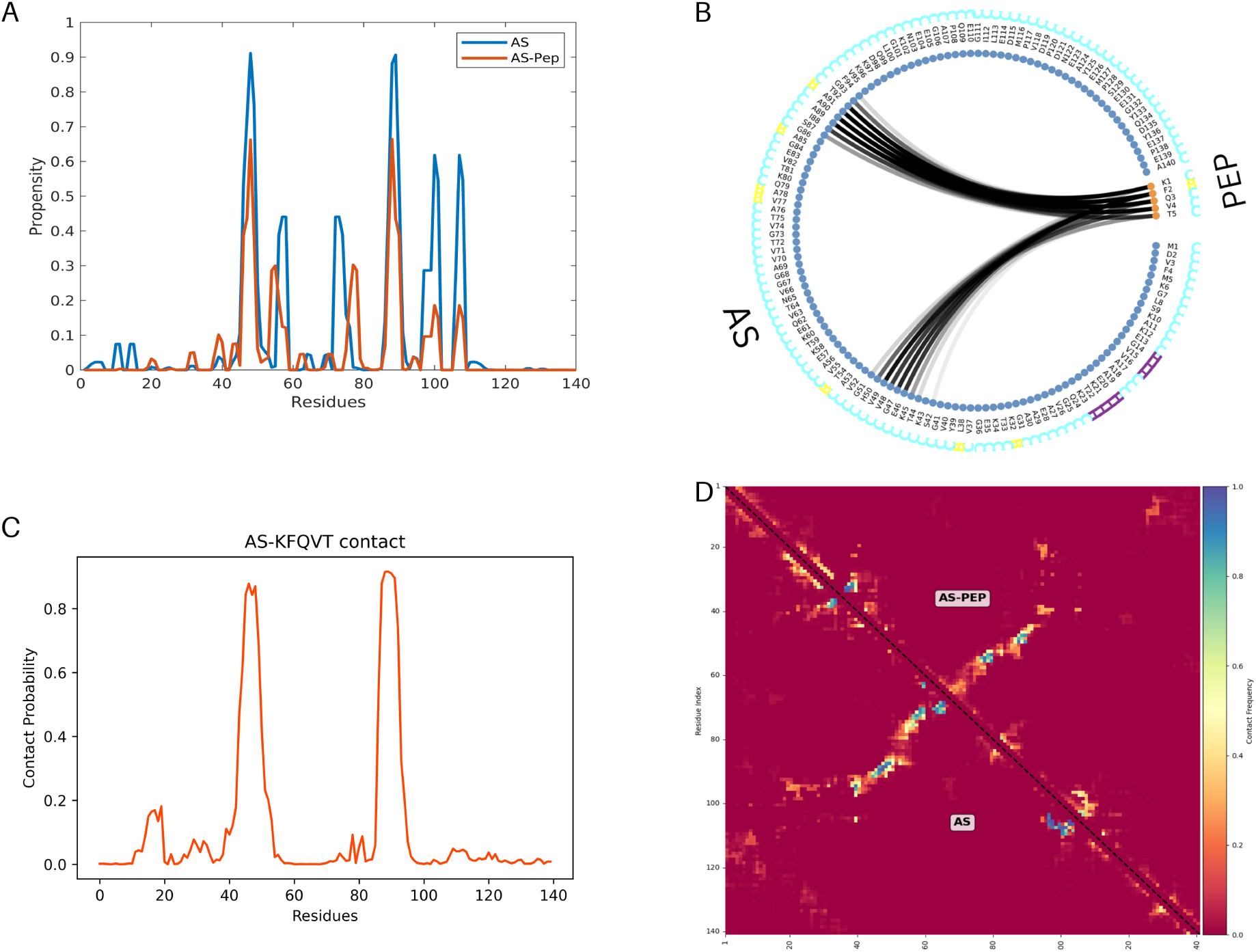
(a) The residue-wise propensity of the *β*-Sheets in the apo and bound states. (b) Binding interaction between *α*-Synuclein and KFQVT peptide. Flare plot representing the protein (AS) residues interacting with the peptide (PEP). (c) Contact probability of the peptide residues with the *α*-Synuclein over the simulation time. Residue-Residue contact map plotted over the period of simulation time for the apo and bound states (d). The graph portrays a noticeable difference in contact between the N and C terminal residues in the bound state. Further, the residues in the NAC region showed reduced contact in the bound state, insinuating the effect of the peptide altering the contact pattern of the *α*-Synuclein system.

The structural changes in the conformations of *α*-Synuclein result from the interaction of the pentapeptide with the NAC domain residues. From the interaction studies, we could infer that the residues A90, A91, and T92 contribute majorly towards the binding with K1, F2, and Q3 of the pentapeptide (Fig. 5B and Fig. 5C). For average statistical read outs, we also plotted the residue-residue contact map (Fig. 5D). The probable contacts formed between the N and the C terminal regions and the NAC region in the apo state are reduced upon interacting with the peptide. Our analysis of residue contacts reveals distinct patterns between the apo state of *α*-Synuclein and its peptide-bound form. In the apo state, we observe high-affinity interactions (probability > 0.8) between residue pairs N65-A69, V70-V63, A69-T64, V71-V63, N65-V71, E46-I88, and V70-Q62. In particular, many of these contacts involve residues within or near the NAC region (residues 61-95), which are crucial for *α*-synuclein aggregation. In contrast, the peptide-bound form demonstrates a different set of high-affinity interactions, including V77-V55, A53-Q79, V77-T54, Q79-V55, A53-A78 and A78-T54. Interestingly, these contacts involve residues both within and outside of the NAC region, suggesting a shift in the protein’s conformational ensemble upon peptide binding. This change in contact patterns indicates that the binding of peptides alters the intramolecular interactions of *α*-Synuclein. Although the apo state demonstrates concentrated interactions within the NAC region, the peptide-bound form exhibits a highly distributed contact pattern. This redistribution of contacts could potentially disrupt the typical aggregation-prone conformations of *α*-Synuclein.

We also computed the vdW contacts within the protein over the simulation time for both apo and bound conformations. Remarkably, we could infer that the residue pairs 60-62, 61-63, 63-70, 63-71, 64-69, 65-68, 65-69, and 97-99 formed significant interaction with the probability above 0.8 in the apo state. Interestingly, all these vdW interactions were lost in the bound state, which clearly shows the interference of the peptide in breaking the intermolecular contacts in the NAC domain leading towards the expanded conformational state of the *α*-Synuclein. For better statistics and to show that expansion seen upon binding was reproducible, we also ran four more replicates each for 1.0 *µ*second (amounting to a total run time of 5.0 *µ*seconds AAMD simulation time for the KFQVT-*α*S complex system).

Taken together, our analyses from the monomeric form of the *α*-Synuclein conformation suggests that the pentapeptide KFQVT pushes the protein towards extended states, reduces the propensity for beta-sheet, weaken the inter-residue interactions at the NAC region and overall provides favorable conditions that could compromise aggregation propensity of *α*-Synuclein chains.

### 2.6 KFQVT affects the stability of ***α***-Synuclein fibrillar structure

To further corroborate the inhibitory activity of the designed peptide against the fibril form *α*-Synuclein, we have utilised the recently resolved cryo-EM model of the pentamer [PDB ID: C6U7] for this study. We used HPEPDOCK tool to dock the peptide KFQVT with the pentameric fibril structure. From the blind docking protocol in HPEPDOCK, we screened out the most energetically favorable docked conformation and used that as our starting structure for the all-atom simulations.

We performed 700 ns AAMD simulations on the pentamer fibrils in the APO and peptide-bound states, respectively (Fig 6A and 6B). We analyzed the secondary structural features from the simulation trajectories since the *β*-sheet propensity is considered to be a hallmark characteristic feature of the fibrillar proteins.^62^ Fig. 6C shows the flare plot with protein residues interactions with the peptide residues. Our analyses show that the percentage of the B-sheet in the *α*-Synuclein apo fibril form (32%) was noticeably reduced upon binding with the peptide (24%) (Fig. 5D). The outcomes from the secondary structure propensity suggest that the presence of peptide has disturbed the *β*-sheet propensity in the *α*-Synuclein fibril form, which transformed into the coil state.

**Figure 6:**
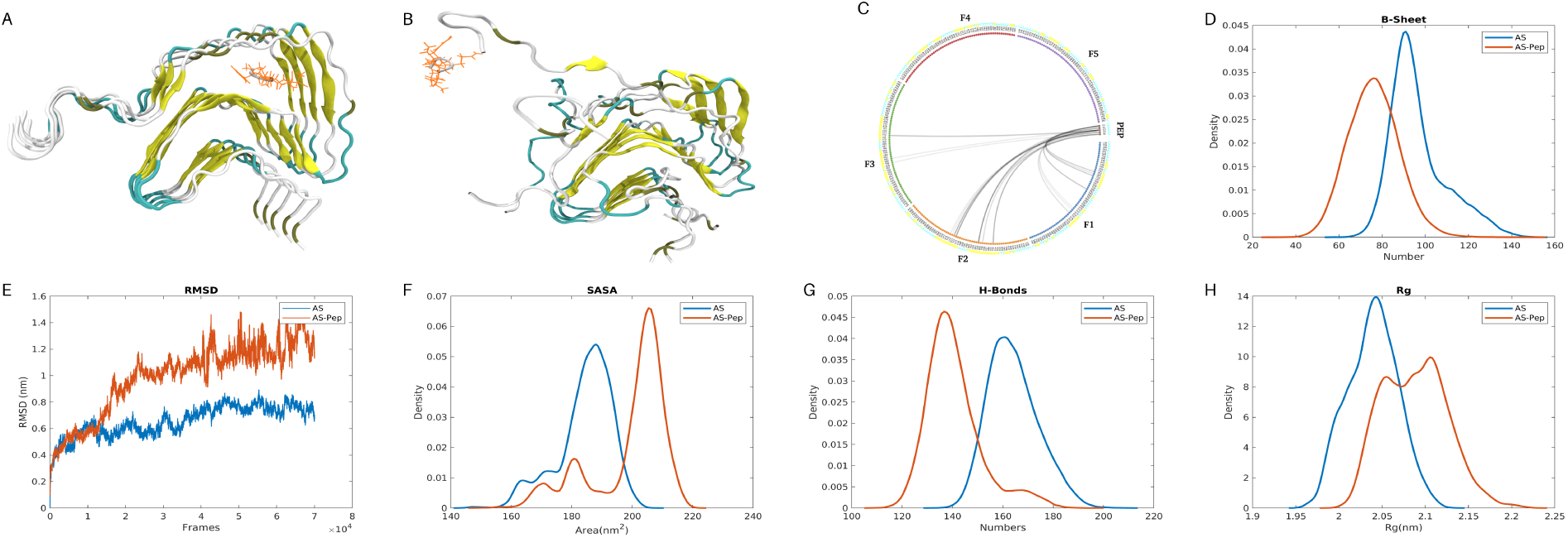
Structural representation and Binding interaction between *α*-Synuclein (fibril) and KFQVT peptide. (a) The *α*-Synuclein (fibril) bound with the designed peptide KFQVT at 0ns. (b) Final snapshot of the fibril complex at 700ns. (c) Flare plot representing the fibril protein (F1-5) residues interacting with the peptide (PEP). Secondary structural propensity of fibril *α*-Synuclein in apo and bound state. (d) *β*-Sheet propensity of the fibril *α*-Synuclein computed for apo and bound state represented in a probability density function. From the graph, we could infer that the binding of the KFQVT peptide has drastically reduced the percentage of the *β*-Sheet in the *α*-Synuclein relative to the apo state. (e) The conformational stability of the fibril system computed using the RMSD parameter denotes a significant difference upon interacting with the peptide. (f) Solvent accessible surface area (SASA) showed that the system is more exposed to solvent in the complex state. (g) Intramolecular hydrogen bonds were noticeably reduced when the peptide interacted with the fibril *α*-Synuclein. (h) The compactness of the protein was found to be exceedingly shifted upon interacting with the peptide.

To further decipher the fibril system’s stability, we computed the structural deviation of the C-*α* atoms in both the APO and bound fibril systems. Fig. 6(E) shows the RMSD of *α*-Synuclein fibril structure in APO and in KFQVT-bound form with the peptide-*α*S showing enhanced instability. Other global readouts such as SASA (Fig. 6(F)), number of hydrogen bonds between the residues of the *α*-Synuclein protein (Fig. 6(G)) and R*_g_* (Fig. 6(H)) shows expanded signatures. Analyses of the trajectory data (see movie files MovieS2) suggests that the presence of charge/polar residues Lysine and Glutamine in peptide could accelerate the preferred binding with E61, T54 and T72 of *α*-Synuclein fibrils, which increases chain flexibility and mediates the loss of conformational stability. The description for each of the movie file for the fibrill system is available in the SI pdf file. The link to the trajectory location can be found in the Github repository.

Furthermore, to authenticate the effect of the designed peptide against the fibril form of *α*-Synuclein, we also carried out simulations with the experimentally studied peptide (KDGIVAGVKA) that was shown to reduce aggregation in *α*-Synuclein.^45^ Remarkably, the outcomes from the control peptide simulations elucidated the inhibitory activity that was similar to what we have observed with the our in silico designed peptide KFQVT (Fig. S1). Results obtained from the control peptide corroborate similar trends in the structural stability, flexibility, compactness, SASA and hydrogen bond propensity relative to the peptide screened using our pipeline. We believe that the designed peptide KFQVT could be taken forward as an aggregation-hindering molecule for *α*-synuclein proteins.

### 2.7 KFQVT lowers the critical temperature of ***α***S condensates

In a recent experimental work using reconstitution and cellular models, Samir Maji and co-workers^18^ demonstrated that liquid–liquid phase separation of *α*-Syn precedes its disease-associated aggregated states. Since the liquid droplet of disordered proteins is possibly a precursor to their pathological fibillar structure,^6,63–65^ it is as such an important therapeutic target. To understand the liquid condensate behaviour of *α*-Synuclein through the computational lens (with and without the screened peptides), we calculated the density of condesnate as as a function of temperature. For our condensate simulations, we used 100 chains of *α*-Synuclein protein and used HPS-Cation coarse-grained force field. ^66^ We use the density data at different temperatures to estimate apparent critical temperature (T*_c_*) of the systems. For methodological details on the high throughput and computationally efficient prescription to crudely estimate T*_c_*, please see the section titled “Estimating apparent critical temperature using PBC density data”.

For the condensate simulations, we use four systems - the first system is amde up of 100 syuclein chains and acts as a control simulation. The other three systems have 100 chains of synuclein and also have 50, 100 and 200 KFQVT peptides, respectively. All four systems were subjected to 5-microsecond simulations with temperatures varying from 380 to 440 K with an increase of 10 K at each simulation. Fig. 7 (a) shows the bulk-phase density for all four systems as a function of temperature. From the data, it is very evident that KFQVT peptides noticeably reduce the density of the condensate especially at higher concentrations. From our condensate simulations, we could infer that the peptides reduced critical temperature *α*-Synuclein condensate (Fig. 7(b)). The apparent T*_c_* drops by as much as 15 K in presence of peptides, which further establishes the effect of peptide on phase separation propensity of syncuelin condesnates.

**Figure 7:**
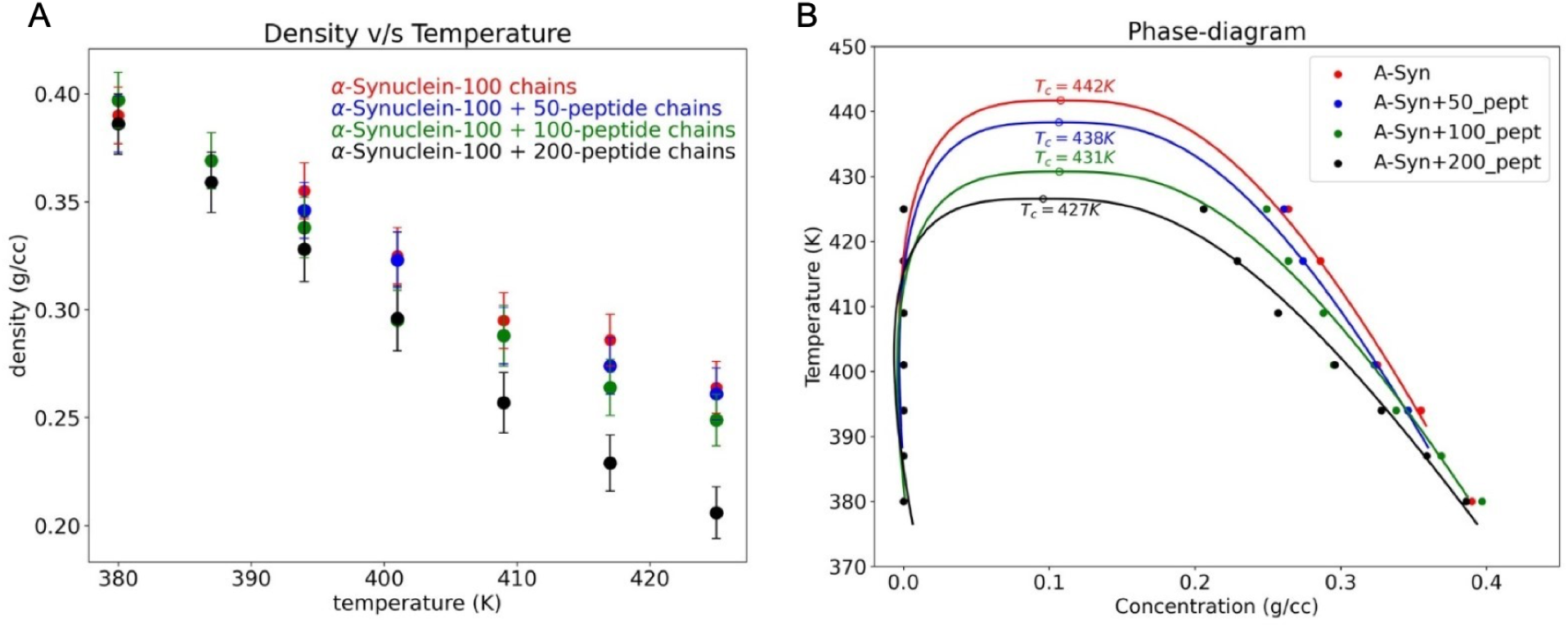
(A) Variation in the density of *α*-Synuclein condensate (with and without peptide) with temperature. (B) Phase-diagram of *α*-Synuclein condensate with and without peptide.

## 3 Conclusions

At the forefront of contemporary drug discovery, there is a growing emphasis on leveraging peptides to address diverse neurological disorders. In this study, we delineate the impact of the *α*-Synuclein-derived peptide (TKEQVTN) by rationally designing the pentapeptide (KFQVT) that intends to impede the formation of *α*-Synuclein aggregates. To gain insights into molecular mechanisms underlying conformational plasticity of *α*-Synuclein in the monomeric, fibrillar, and condensate forms with the presence of the derived peptide inhibitor, we employed molecular dynamics simulations at both atomistic and coarse-grained resolutions. From the atomistic *µ*seconds simulations, we have characterized the binding of the KFQVT peptide to the monomeric *α*-Synuclein that revealed the unique binding interactions. In particular, the KFQVT binds near the NAC region of the *α*-Synuclein and induces changes in the structural compactness, solvent accessibility and secondary structure propensity in the bound form. The results show that the interactions are driven by entropic factors, which could arise from the increase in the conformational entropy of the *α*-Synuclein through the entropic expansion mechanism.

Further, we investigated the effect of the peptide against the fibrillar form of the *α*-Synuclein at the atomic level. Results from our simulations showed that upon binding, the formation of *β*-sheets was profoundly reduced in *α*-Synuclein relative to that of apo-form with a noticeable difference in their propensity. Moreover, the preferential binding of KFQVT altered the contact pattern, leading to increased structural compactness and flexibility. Further, these structural changes led to more solvent exposure and loss in internal hydrogen bonds, leading to overall change in the conformational stability of the system. Moreover, our study showed that the designed peptide exhibited similar inhibitory activity relative to the experimentally studied peptide in disturbing the *α*-Synuclein aggregation. While our results align considerably with experimental studies, further experimental validation is necessary to establish the effectiveness of the peptide.

### Scope and limitations

While our multiscale simulation pipeline offers a cost-effective platform for initial screening, it is essential to acknowledge several limitations. Molecular dynamics simulations, although providing valuable mechanistic insights, may not fully capture the intricate complexities of the long-term dynamics of aggregation processes within a cellular environment. Moreover, the in vivo stability, bioavailability, and potential off-target effects of the modified peptide remain to be experimentally determined. To establish the therapeutic potential of these peptides, rigorous experimental validation in cellular and animal models is imperative to corroborate the computational predictions. Despite these limitations, our findings establish a foundation for further investigations in to peptide-based inhibitors of *α*-Synuclein aggregation and underscore the potential of computational methodologies in drug discovery for neurodegenerative diseases.

## 4 Computational Methods and Systems

### 4.1 Docking studies on monomeric system

Once the structural conformation of the peptide library was built, we started our initial study with protein-peptide docking. We used PatchDock, a rigid-body docking algorithm based on shape complementarity, to generate a pool of potential binding poses with all 14 full-length *α*-Synuclein conformations obtained from the clustering data. Relative binding energy values for these complexes were calculated using Prodigy^53^ to assess if the predicted structures were energetically favorable. Subsequently, the top-scoring poses were subjected to refinement using HPEPDOCK, which is a hierarchical algorithm that incorporates flexible peptide docking. This combined approach leverages the efficiency of PatchDock for rapid identification of potential binding modes with the precision of HPEPDOCK for accurate pose prediction and scoring. We chose the top binding conformation as input for our molecular simulation studies.

### 4.2 Redesigning the final peptide

To enhance peptide binding affinity, the CSM-PPI web server was employed. The mCSM-PPI web server uses graph-based structural signatures to represent the wild-type residue environment, which are then used as input features for machine learning models. The server integrates these structural signatures with evolutionary information and interatomic interactions to provide quantitative predictions of changes in binding affinity upon mutation. CSM-PPI algorithm identifies key residues critical for peptide-*α*-Synuclein binding and helps us with the amino acid substitutions that could potentially augument binding affinity while preserving peptide stability. This computational approach offers a streamlined iterative method for an optimized protein for a given complex.

### 4.3 Peptides with monomeric ***α***Synuclein: All-atom simulations

GROMACS was used for the all-atom molecular dynamics simulations with the amber99sb-disp force field in combination with the amber99sb-disp water models. ^49^ This combination of force fields was shown to capture the conformation ensembles relative to the experimental measurements for both folded and intrinsically disordered proteins. *α*-Synuclein chain was kept in the rectangular box with periodic boundary conditions, solvated and neutralized by adding counter ions (NA+ and CL-) to the concentration of 150 mM. The systems were energy-minimized using the steepest descent algorithm and equilibrated for 100 ps in the canonical and isothermal isobaric ensemble at 310 K and 1 atm, respectively. The production run was started using the well-equilibrated system with a v-rescale thermostat and Parrinello-Rahman barostat. ^67^ Water molecules are kept rigid using the SETTLE algorithm. Non-water bonds involving hydrogen atoms were restricted using the LINCS algorithm, providing a time-step of 2 fs for integrating the equations of motion. Particle Mesh Ewald(PME) approach was used for long-range interactions. ^68^ No restraints were kept on the peptide to the binding site, allowing it to move around the system freely. We followed two independent simulations for all the systems. We have analysed the trajectories using the GROMACS-built tools such as g_rmsf, g_sasa, g_hbond, and g_gyrate for calculating residual flexibility, solvent accessibility, hydrogen bonds and radius of gyration. We also compute the secondary structure propensities using the do_dssp tool. All the graphs were plotted using the MATLAB. Chimera was used for the visualisation.

### 4.4 Peptides with Fibril system: AAMD simulations

We have chosen the pentamer of Cyro-EM structure deposited with PDB id: 6CU7 to study the effect of the designed peptide against the fibrillar form of the *α*-Synuclein. Structurally, the 6CU7 has been resolved with 60 residues from L38-K97.^13^ The selected structure was subjected to molecular docking experiments with our designed peptide using the HPEPDOCK program. The complex with the highest binding affinity was selected as the initial structure for subsequent all-atom molecular dynamics simulations using GROMACS. The apo state served as the control for both monomeric and fibril form systems. The simulation systems were constructed using periodic boundary conditions with cubic box dimensions of 88 Å × 88 Å × 88 Å for the fibril system. System neutralization was achieved through the addition of Na+ and Cl-counterions to maintain electrostatic neutrality. Long-range electrostatic interactions were handled using the Particle Mesh Ewald (PME) method, with Coulomb cut-off radii implemented. Prior to production runs, each system was energy minimized and equilibrated to ensure proper relaxation of the systems. The simulations were carried out using 700 ns for each of the systems. Force field parameters and simulation conditions were consistently maintained across all atomic-level simulations to ensure methodological uniformity. Detailed specifications regarding system size, dimensions, and simulation duration are systematically presented in Table 1.

**Table 1:** List of simulated all atom systems.

| Systems | Box Dimensions<br>(Åx Åx Å) | Number<br>of atoms | Simulation<br>time (ns) |
| --- | --- | --- | --- |
| α-Syn monomer | 109 x 109 x 160 | 248541 | 5000 |
| α-Syn monomer<br>with peptide | 109 x 109 x 160 | 248496 | 5000 |
| α-Syn fibril | 88 x 88 x 88 | 88715 | 700 |
| α-Syn fibril – pep-<br>tide | 88 x 88 x 88 | 88724 | 700 |
| α-Syn fibril- con-<br>trol peptide | 88 x 88 x 88 | 88700 | 700 |

### 4.5 CG simulations using HPS-Cation-***π*** and apparent critical temperature calculations

To study the molecular interactions governing the biomolecular condensate of *α*-Synuclein, we used the LAMMPS simulation package to perform coarse-grained simulations with 100 *α*-Synuclein chains and different titrations of KFQVT. To model the core region of the condensate, we choose PBC box condensate simulation rather than the conventional direct coexistence slab-based condensate simulation method (see Fig. 8(A,B). We use the HPS-Cation-pi coarse-grained force field. ^66,69^ We have taken hundreds of chains of *α*-synuclein with and without peptides in a cubic box and simulated them with NPT ensemble. The timestep for the Verlet integration of the equations of motion was chosen to be of 10 fs. NPT simulations performed for pure bulk protein liquids were carried out at p=1 bar using a Nosé-Hoover barostat for 1 *µs* simulation. We first establish that the core density of the slab system (Fig. 8C) was equal to the PBC density reported from our simulations. Fig. 8(E) shows the core density at different temperatures for the 100-chain *α*-Synuclein system. For a high throughput calculations of “apparent” critical temperatures and compare different condensate systems, we assume the density of the dilute phase to be zero and fit the densities data to extract the co-existence curve (Fig. 8(F)). The following two equations were solved simultaneously to obtain the critical temperature T*_c_* and density *ρ_c_*. Unlike in the case of 3D Ising model and Lennard Jones system, *β* is set to be a free parameter in the fits. The fitting is iterated for different values of fit parameters A and Δ*ρ_o_* to obtain T*_c_* and density *ρ_c_* values corresponding to least error. The final reported values correspond to best fits of the binodals.

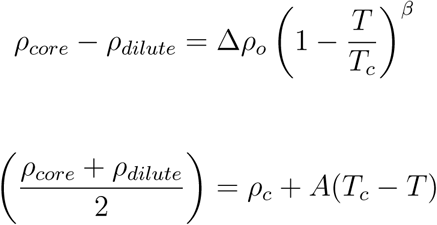

**Figure 8:**
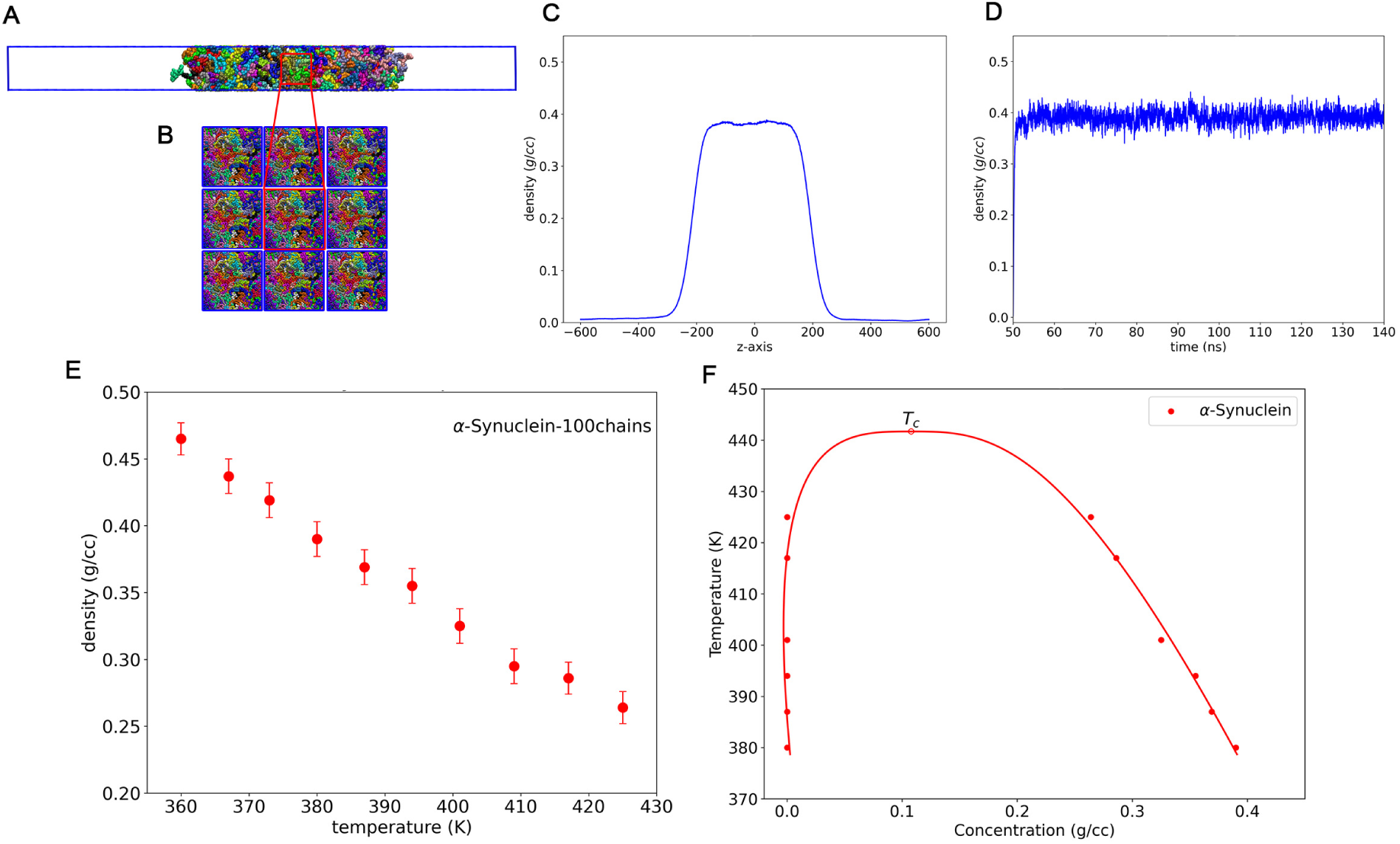
(A) The direct coexistence slab simulation, which consists of two phases of condensate, dilute and dense phase. (B) PBC box simulation, which only has a dense phase of condensate. (C) Density profile of slab simulation. (D) density convergence of PBC box condensate simulation. (E) variation in the density of pure *α*-Synuclein 100 chains with temperature. (F) Phase diagram of pure *α*-Synuclein.

All analyses were performed using the MDAnalysis package and in-house Python scripts that is made available in our Github repository. Input files for the CGMD simulations are provided in the Github repository and links to all trajectories are also available there.

## Acknowledgement

ES thanks the Department of Biotechnology (DBT), Government of India, and IISc for the Institute of Eminence (IoE) postdoctoral fellowship and SP thanks the Ministry of Education, Government of India, for the graduate fellowship. AS acknowledges the financial support from the Indian Institute of Science (IISc) and the high-performance computing facility “Beagle” that was set up from grants by the erstwhile IISc-DBT partnership programme. AS thanks the DST for the National Supercomputing Mission grants (DST/NSM/R&D-HPC-Applications/2021/03.10, DST/NSM/R&D-HPC-Applications/Extension Grant/2023/27). AS also acknowledges the FIST program sponsored by the Department of Science and Technology, India that supports the MBU infrastructure. AS would also like to thank the Teams Science Grant from the DBT-Wellcome Trust India Alliance (Grant number: IA/TSG/21/1/600245). AS also thanks the DBT National Network Project (NNP) grant (BT/PR40323/BTIS/137/78/2023) and the Matrics grants (MTR/2023/001040) from the Science and Engineering Board (SERB), India.

## Author contributions

AS conceptualized the project. ES and AS designed the research. ES and SP performed the research and analyzed the data. ES worked on peptide design, molecular docking, and worked needed to design of final candidate peptide. ES also carried out AAMD simulations on single chain *α*-Synuclein single molecule and fibril systems. SP worked on coarse-grained modeling and simulations of *α*-Synuclein condensate and calculations relayed to density and apparent critical temperatures. AS supervised the study. ES prepared the first draft of the paper with inputs from SP. AS polished the draft.

## 6 Data availability statement

Our code, models, and curated datasets are publicly available at the laboratory GitHub repository https://github.com/codesrivastavalab/synuclein-KFQVT.

## Supporting Information

### List of movie files from MD trajectories

**Table S1:** The table represents the top peptides in each library having better affinity with the *α*-Synuclein conformations computed via prodigy. The top peptides were further subjected to HPEPDOCK for accurate prediction of the protein peptide pose prediction with affinity score. Library (AA): Libraries build accoringly for the length of peptide sequences. PEP-SEQ: Sequence of the peptide; Peptide_no-AS_struct: Peptide which was obtained from the *α*-Synuclein structure. The results of the binding affinity are reported in Kcal/mol.

| Lib | PEP-SEQ | Peptide# | Patchdock Score | Hpepdock Score | Residence Time |
| --- | --- | --- | --- | --- | --- |
| 3 | QKT | Pep79-41 | -7.1 | -105.126 | 200 ns |
| 4 | QVTN | Pep62-28 | -9 | -82.399 | 100 ns |
| 5 | EQVTN | Pep61-41 | -8.5 | -90.045 | 100 ns |
| 6 | KEQVTN | Pep60-28 | -9.6 | -105.228 | 100 ns |
| 7 | TKEQVTN | Pep59-1 | -10.3 | -112.337 | 350 ns |
| 8 | KKDQLGKN | Pep96-1 | -10.6 | -108.899 | 200 ns |
| 9 | KTKEQVTNV | Pep58-41 | -11 | -146.448 | 290 ns |

**Table S2:** The table represents the redesigned peptides having better affinity with the *α*-Synuclein conformations computed via CSM-PPI. Mutations at position 2 (W, P, F, Y) of the peptide KEQVT, predicted to enhance *α*-Synuclein binding affinity by CSM-PPI, were further evaluated using HPEPDOCK and HAWKDOCK. HPEPDOCK conducts flexible peptide-protein docking, while HAWKDOCK employs short MD simulations and MMGBSA calculations for energy estimation. PEP-SEQ-Sequence of the peptide; The results of the binding affinity are reported in Kcal/mol.

| PEP-SEQ | Hpepdock Score | MMGBSA Score |
| --- | --- | --- |
| KEQVT | -97.427 | -32.97 |
| KWQVT | -129.64 | -42.03 |
| KPQVT | -116.808 | -31.96 |
| KFQVT | -132.624 | -43.39 |
| KYQVT | -123.346 | -33.54 |

**Fig. S1:**
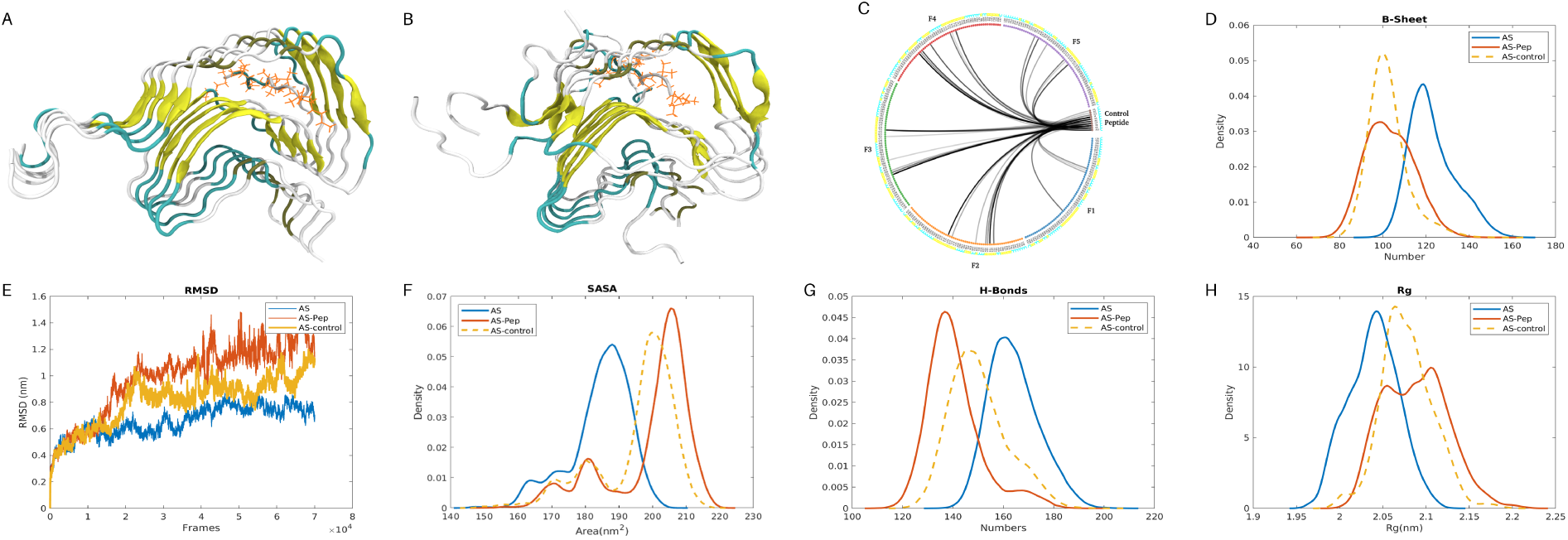
Comparative structural analysis of the apo, KFQVT and control peptide bound fibril states, respectively.

1. **movieS1a.mpeg**: Movie file of KFQVT peptide bound *α*-Synuclein chain in solution state from 1 microsecond all-atom trajectory data (replicate #1).
2. **movieS1b.mpeg**: Movie file of KFQVT peptide bound *α*-Synuclein chain in solution state from 1 microsecond all-atom trajectory data (replicate #2).
3. **movieS1c.mpeg**: Movie file of KFQVT peptide bound *α*-Synuclein chain in solution state from 1 microsecond all-atom trajectory data (replicate #3).
4. **movieS1d.mpeg**: Movie file of one of the instances of full-length *α*-Synuclein chain in solution state from 1 microsecond all-atom trajectory data.
5. **movieS2a.mpeg**: Movie file of KFQVT peptide bound *α*-Synuclein fibrill systems from 1 microsecond all-atom trajectory data.
6. **movieS2b.mpeg**: Movie file of previously experimentally studied peptide (KDGI-VAGVKA)^1^ bound *α*-Synuclein fibrill systems from 1 microsecond all-atom trajectory data (for comparative structure study shown in Fig. S1).
7. **movieS2c.mpeg**: Movie file of *α*-Synuclein fibrill systems from 1 microsecond all-atom trajectory data (control simulation).

